# RNA Polymerase II coordinates histone deacetylation at active promoters

**DOI:** 10.1101/2024.09.17.613553

**Authors:** Jackson A. Hoffman, Kevin W. Trotter, Trevor K. Archer

## Abstract

Nucleosomes at actively transcribed promoters have specific histone post-transcriptional modifications and histone variants. These features are thought to contribute to the formation and maintenance of a permissive chromatin environment. Recent reports have drawn conflicting conclusions about whether these histone modifications depend on transcription. We used triptolide to inhibit transcription initiation and degrade RNA Polymerase II and interrogated the effect on histone modifications. Transcription initiation was dispensable for *de novo* and steady-state histone acetylation at transcription start sites (TSSs) and enhancers. However, at steady state, blocking transcription initiation increased the levels of histone acetylation and H2AZ incorporation at active TSSs. These results demonstrate that deposition of specific histone modifications at TSSs is not dependent on transcription and that transcription limits the maintenance of these marks.

## Main Text

In transcribing the genome, the RNA polymerase II (RNAP2) complex traverses variably compacted chromatin in the form of nucleosomes and higher-order structures. Multiple protein complexes function to remodel chromatin and modify nucleosomes to permit the formation and maintenance of different genomic environments. For instance, actively transcribed regions of the genome coincide with nucleosomes containing the H2AZ histone variant and acetylated histones(*1*). However, the precise functional relationship between transcription and these active chromatin modifications is ill-defined. Recent work has concluded that several TSS-associated histone modifications were dependent on transcription(*2, 3*). Conversely, other studies have demonstrated that active marks persist upon inhibition of transcription(*4, 5*). Moreover, inhibition of the histone acetyltransferases (HATs) and histone methyltransferases (HMTs) that deposit active marks can prevent recruitment of the pre-initiation complex, alter RNAP2 occupancy, and/or block transcription(*6-10*). Thus, despite their well-established association, the relationship between active transcription and histone modifications presents a causality dilemma(*11*).

### Transcription is dispensable for steady-state and *de novo* H3K27 acetylation

Stimuli-driven responses present an attractive model for resolving this dilemma, as exogenous stimuli can induce rapid changes in both transcription and the distribution of histone modifications. For instance, the synthetic glucocorticoid dexamethasone (dex) induces widespread changes in RNAP2 binding, transcription, histone acetylation, and chromatin accessibility within one hour of treatment in human T47D-derived A1-2 breast cancer cells(*12*). To interrogate the hierarchy of events in this response, we sought to block transcription and examine the effects on steady-state and hormone-induced histone modifications. To do so, we used triptolide, a natural product that blocks transcription initiation by inhibiting the ATPase activity of the XPB subunit of TFIIH, resulting in CDK7-dependent hyperphosphorylation and degradation of RNAP2(*13*). We pre-treated cells with triptolide for 1 hour to induce RNAP2 degradation followed by co-treatment with triptolide and dex for 1 hour. This treatment regimen resulted in degradation of RNAP2 and prevented dex-induced recruitment of RNAP2 to Glucocorticoid Receptor (GR) target genes such as ZBTB16 and GLUL (Figure 1A-D, S1).

**Figure 1.**
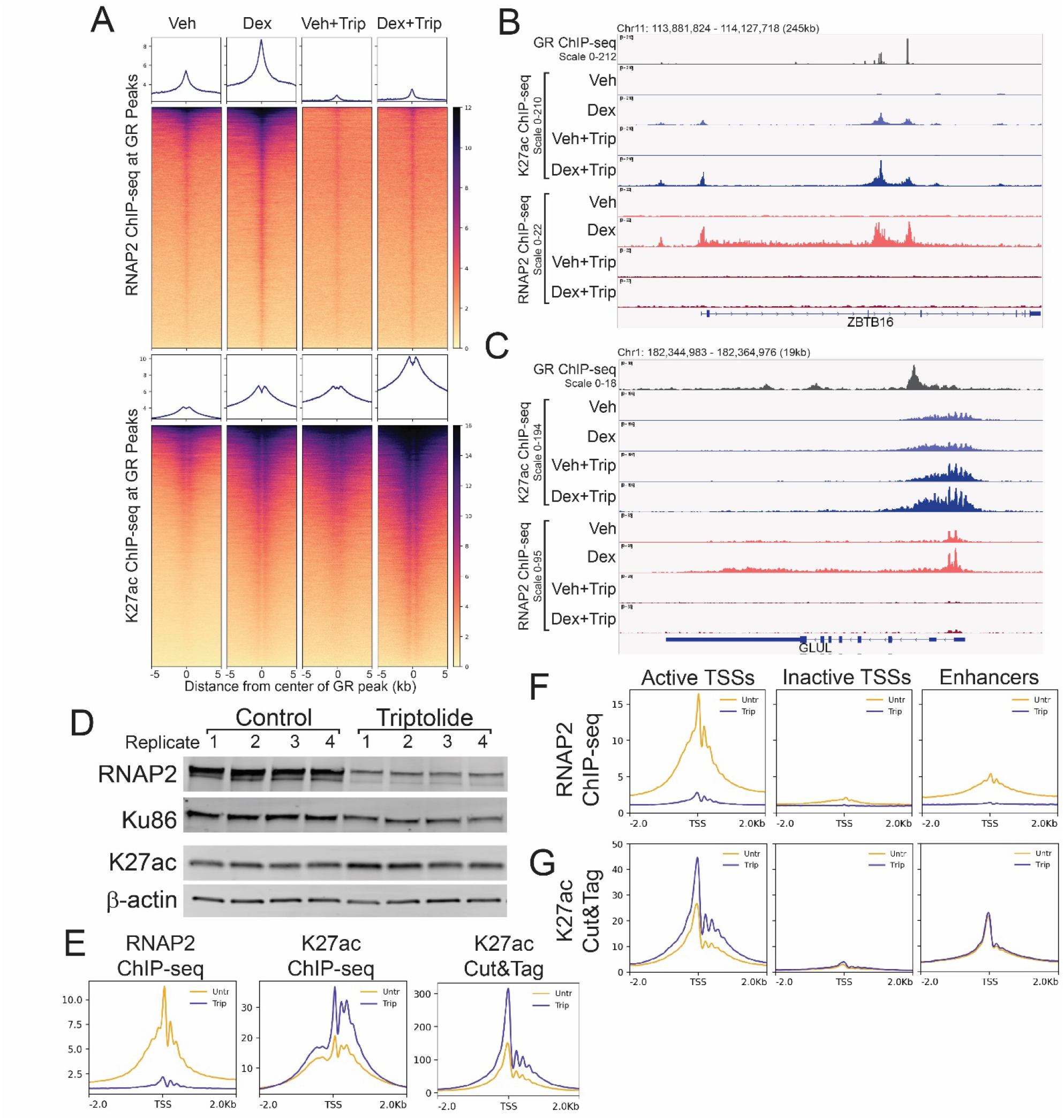
Transcription is dispensable for steady-state and *de novo* H3K27 acetylation. A) Differential heatmap of K27ac ChIP-seq signal (log2 Dex/Veh) over Class III GR peaks +/-2 hours triptolide. B) Browser image of K27ac ChIP-seq and RNAP2 ChIP-seq over ZBTB16 gene. C) Browser image of K27ac ChIP-seq and RNAP2 ChIP-seq over GLUL gene. D) Western blot of whole cell lysates from 4 biological replicates each of A1-2 cells +/-2 hours triptolide. E) Meta-profiles of representative ChIP-seq and Cut&Tag replicates over Refseq TSSs. F) Meta-profiles of representative RNAP2 ChIP-seq +/-2 hours triptolide over Start-seq defined active, inactive, and enhancer TSSs. G) Meta-profiles of representative K27ac Cut&Tag +/-2 hours triptolide over Start-seq defined active, inactive, and enhancer TSSs.

GR binds to ~30,000 sites in the A1-2 genome after 1 hour of dex treatment. Hormone-dependent recruitment of RNAP2 and acetylation of histone H3 lysine 27 acetylation (K27ac) is evident over these regions. (Figure 1A). While RNAP2 recruitment was mostly abolished, triptolide treatment did not prevent the hormone-induced increase in K27ac at GR peaks (Figure 1A). Rather, K27ac levels were increased in both vehicle and dex conditions with triptolide (Figure 1A). At ZBTB16, *de novo*, dex-dependent K27ac was observed at 2 intragenic GR peaks and at the TSS where GR binding was not detected (Figure 1B). At all 3 sites, triptolide treatment resulted in even greater K27ac upon dex treatment (Figure 1B). Thus, dex-dependent acetylation of K27 occurred in the absence RNAP2 recruitment and transcription initiation.

At GLUL, GR binding downstream of the TSS does not trigger much change in K27ac but does result in increased RNAP2 at the TSS and in the gene body (Figure 1C). Triptolide treatment increased K27ac over the TSS independently of dex, suggesting that triptolide induced an increase in steady-state K27ac (Figure 1C). Indeed, triptolide treatment resulted in a small increase in the overall levels of K27ac detected in whole cell lysates and enhanced K27ac ChIP-seq signal over TSSs (Figure 1D-E, S1). K27ac Cut&Tag confirmed that triptolide treatment resulted in enhanced K27ac signal over TSSs (Figure 1E).

We previously used Start-seq to identify sites of RNAP2 transcription initiation in A1-2 cells(*12, 14*). This allowed us to classify three sets of TSSs: I) active gene TSSs associated with Start-seq signal, II) non-transcribed or inactive gene TSSs, and III) actively transcribed enhancers. We used these TSSs subsets to determine whether the increased levels of K27ac following inhibition of initiation were restricted to TSSs where RNAP2 was active. While triptolide induced RNAP2 loss at both active TSSs and enhancers, the increased levels of K27ac were only observed at active gene TSSs (Figure 1F-G). This increase in K27ac levels at active gene TSSs was validated over 5 independent biological replicates of ChIP-seq and 8 independent biological replicates of Cut&Tag (Figure S2). Thus, K27ac persisted at increased levels at active gene TSSs following inhibition of transcription initiation.

### Transcription initiation suppresses acetylation and H2AZ incorporation at active TSSs

To determine whether this increase was unique to K27ac, we examined additional active chromatin modifications by Cut&Tag. Inhibition of transcription initiation resulted in increased levels of acetylation at H3K9, K3K14, H3K18, H4K5, and H4K12 at active TSSs without noticeable change in bulk levels (Figure 2A-E, S3). However, tri-methylation of H3K4 (H3K4me3) at active TSSs was unaffected by inhibition of initiation (Figure 2F, S3). Similarly, the H3K27me3 repressive mark was also unaffected (Figure 2G). Thus, both acetylation and methylation of histones at TSSs persisted, and the levels of histone acetylation were increased in the absence of transcription initiation.

**Figure 2.**
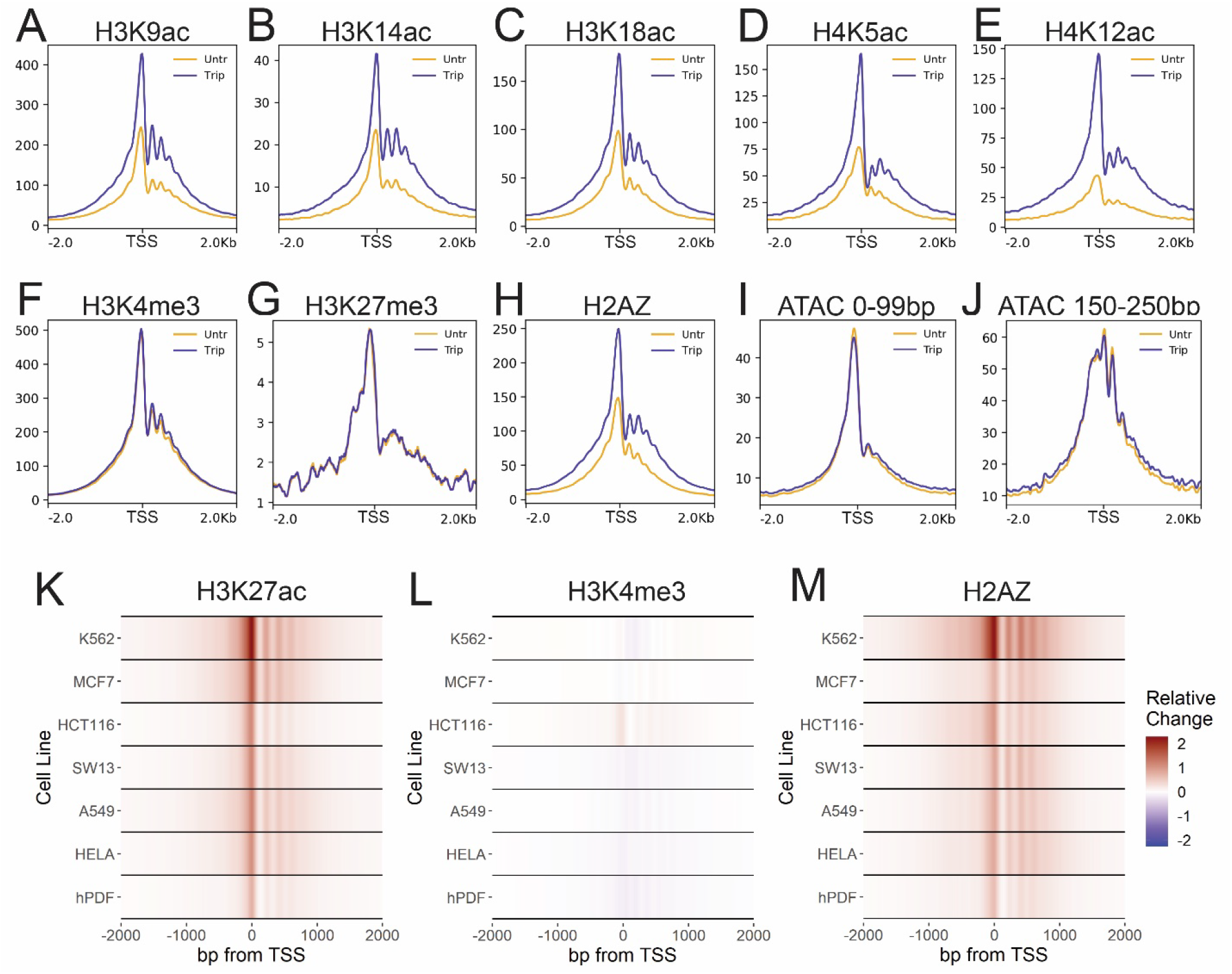
Transcription initiation suppresses acetylation and H2AZ incorporation at active TSSs. A-J) Meta-profiles of representative Cut&Tag replicates +/-2 hours triptolide over active TSSs. K-M) Heatmaps depicting relative change ((Triptolide – control) / max(control)) in Cut&Tag signal over Refseq TSSs in indicated cell lines.

Active TSSs are also marked by the replacement of histone H2A with the histone variant H2AZ. We also observed enhanced levels of the H2AZ incorporation at active TSSs after inhibition of transcription initiation (Figure 2H). Although H3K4me3 was unchanged, the observed increase in acetylation and H2AZ incorporation could be caused by a change in the nucleosome occupancy following triptolide treatment. To test this, we performed ATAC-seq to examine chromatin accessibility at active TSSs. Inhibition of initiation had little effect on the profile of either short, nucleosome-free fragments or nucleosome-size fragments (Figure 2I-J). Therefore, the inhibition of transcription initiation enhanced both histone acetylation and H2AZ incorporation on nucleosomes at active TSSs.

To further validate these findings, we examined the effect of transcription inhibition on K27ac, H3K4me3, and H2AZ enrichment in additional cell lines. As in A1-2 cells, we observed increased K27ac and H2AZ at TSSs following triptolide treatment in human lymphoma (K562), breast (MCF7), colon (HCT116), adrenal gland (SW13), lung (A549), and cervical (HeLa) cancer cell lines and in primary dermal fibroblasts (hPDF) (Figure 2K, M). H3K4me3 remained largely unchanged after triptolide treatment in each of these cell lines, apart from HCT116 cells which exhibited a marginally increased level (Figure 2L). Thus, the effects of inhibiting transcription initiation were highly reproducible and largely independent of cell line and tissue-of-origin.

Previously, inhibition of transcription with triptolide has been shown to trigger a global loss of K27ac and H3K4me3 in K562 cells(*3*). The ChIP-seq experiments performed in this study followed a different protocol with milder cross-linking and MNase fragmentation (*3*). To determine whether the discrepancy in findings was due to technical differences, we repeated our ChIP-seq experiment using these conditions and reagents. With this alternative protocol, we observed similar profiles of increased K27ac and H2AZ along with unchanged H3K4me3 at TSSs following triptolide treatment (Figure S4). As such, our findings are consistent across distinct ChIP-seq protocols as well as Cut&Tag.

As triptolide rapidly blocks transcription initiation, we sought to profile the changes in acetylation and H2AZ incorporation over a time course of treatment. Increased levels of K27ac and H2AZ were observed within 15 minutes of triptolide treatment, with a further increase at 30 minutes (Figure S5). Taken together, these experiments demonstrated that blocking transcription initiation rapidly resulted in enhanced acetylation and H2AZ incorporation at TSSs.

### RNAP2 degradation reproduces the effect of inhibiting transcription inhibition

To investigate whether the enhanced acetylation and H2AZ incorporation was caused by the loss of transcription initiation or RNAP2 degradation, we utilized an alternative method of inhibiting transcription. Flavopiridol inhibits CDK9 and prevents phosphorylation of NELF and DSIF to block pause release(*15*). Treatment with flavopiridol for 2 hours stalled RNAP2 at TSSs and blocked transcription elongation (Figure 3A). This resulted in a marginal increase in K27ac, unchanged H3K4me3, and unchanged H2AZ at active TSSs (Figure 3B-D). Thus, blocking transcription by inhibiting pause release was not sufficient to trigger the increased histone acetylation observed after blocking transcription initiation.

**Figure 3.**
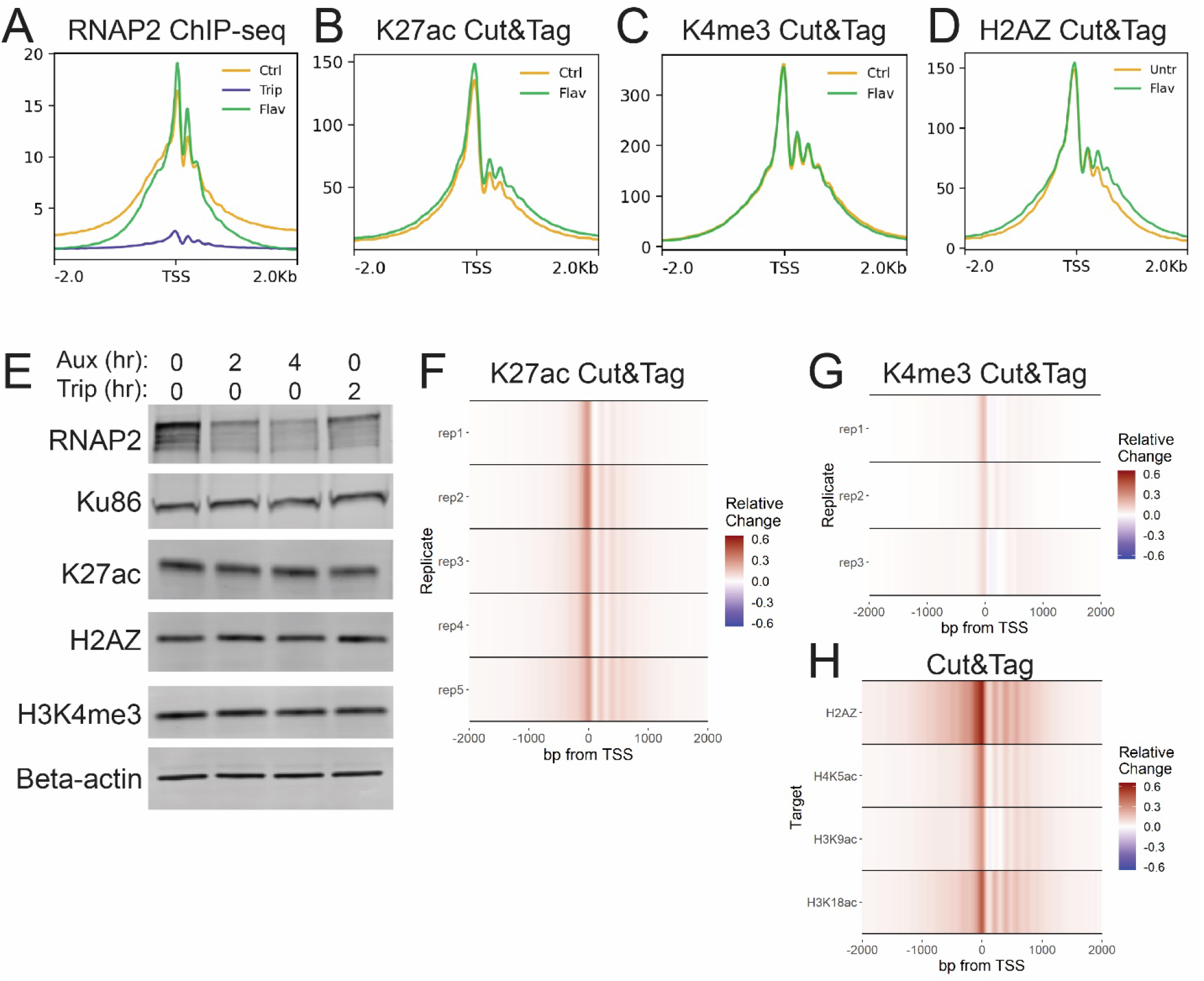
RNAP2 degradation reproduces the effect of inhibiting transcription inhibition. A-D) Meta-profiles of ChIP-seq and Cut&Tag signal +/-2 hours flavopiridol over active TSSs. E) Western blots of whole cell lysate from HCT116-mAID-POLR2A cells +/-Auxin or triptolide. F-H) Heatmaps depicting relative change ((auxin – control) / max(control)) in Cut&Tag signal for indicated marks.

We next sought to determine whether RNAP2 degradation was sufficient to replicate the effect of triptolide on histone acetylation and H2AZ incorporation. To do so, we utilized an auxin-inducible degron (AID) system in HCT116 cells in which RPB1 is bi-allelically tagged with a mini-AID and mClover cassette(*16*). These cells also harbor a doxycycline-inducible cassette for expression of the Oryza sativa auxin-binding protein TIR1 (osTIR). Following 48 hours of dox induction of osTIR expression, rapid degradation of RNAP2 can be triggered by treatment with auxin. 2 hours of auxin treatment was sufficient to reduce RNAP2 similarly to 2 hours of triptolide treatment without noticeable change in the bulk levels of histone modifications (Figure 3E, S6). This degradation of RNAP2 resulted in reproducibly increased K27ac at active TSSs, albeit a lower level than observed with triptolide (Figure 3F). A milder increase was observed in H3K4me3, similar to that caused by triptolide in HCT116 cells (Figure 3G). Levels of H2AZ incorporation, H3K9ac, H3K18ac, and H4K5ac were also increased after AID degradation of RNAP2 (Figure 3H). This revealed that depletion of RNAP2 without inhibition of transcriptional activity could also trigger enhanced histone acetylation and H2AZ incorporation at TSSs.

### HDAC inhibition masks the effect of inhibiting transcription initiation

Histone acetylation is dynamically regulated by the activity of histone acetyltransferases (HATs) and histone deacetylases (HDACs), which respectively catalyze the deposition and removal of acetyl marks. We next sought to determine whether the increased acetylation caused by blocking transcription initiation was the result of increased HAT activity or a loss of HDAC activity. To test HAT-dependence, we used the P300 inhibitor A485(*17*). P300 is the predominant HAT responsible for K27ac at TSSs(*18*). Treatment with A485 diminished the levels of K27ac detected in cell lysates and at TSSs but had little effect on H3K4me3 or H2AZ (Figure 4A-D, S7). Despite the reduced overall levels of K27ac, A485 did not prevent the triptolide-induced increase in K27ac or H2AZ at active TSSs. Thus, the gained acetylation at active TSSs was not dependent on P300 activity.

**Figure 4.**
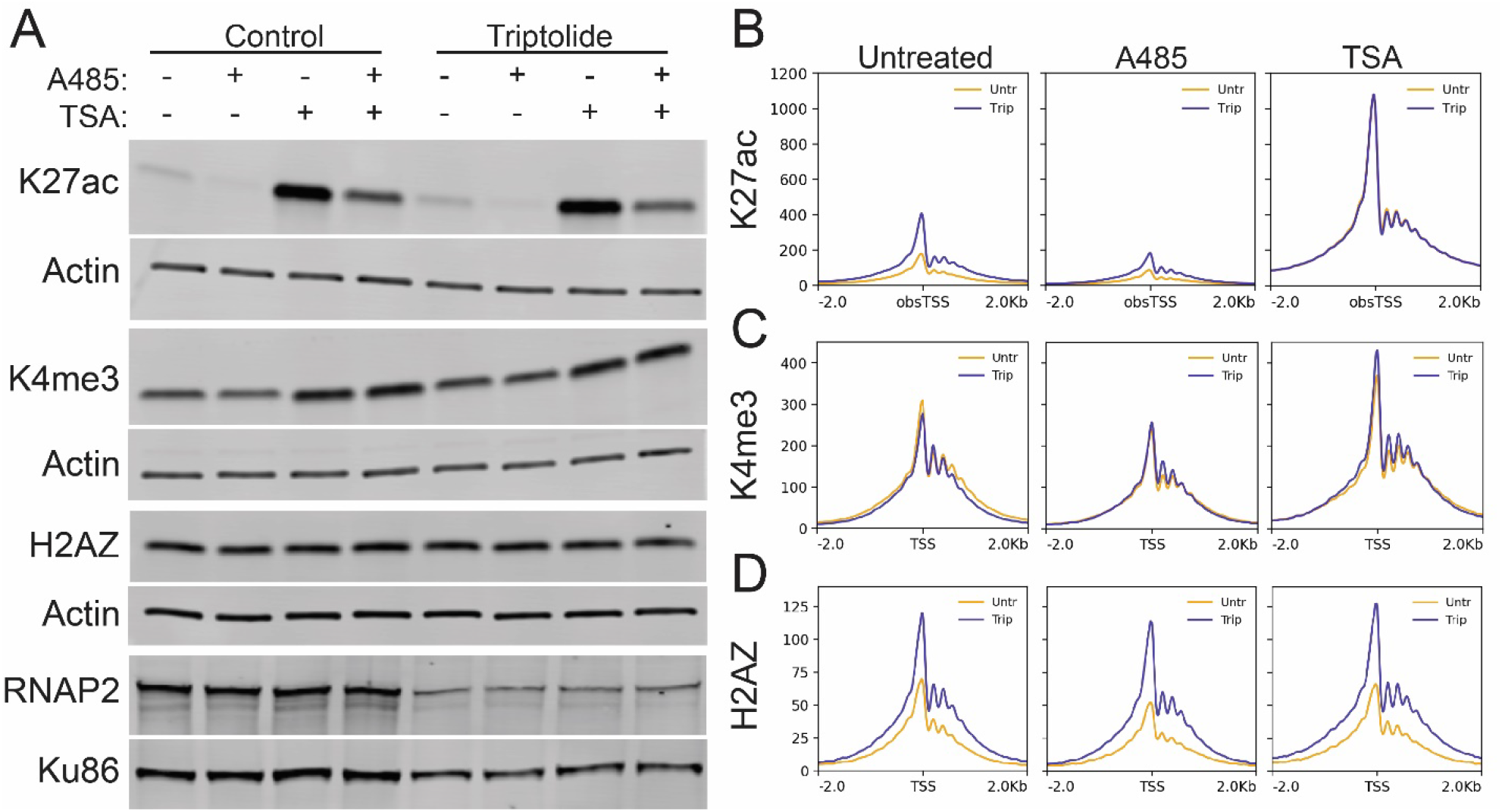
HDAC inhibition masks the effect of inhibiting transcription initiation. A) Western blots of whole cell lysates from A1-2 cells treated +/-triptolide, A485, and TSA. B-D) Meta-profiles of Cut&Tag signal in A1-2 cells treated +/-triptolide, A485, and TSA.

To test whether a loss of HDAC activity was responsible for the gained acetylation, we treated cells with trichostatin A (TSA), an inhibitor of class I and class II HDACs(*19*). TSA treatment markedly increased the levels of K27ac in cell lysates and at active TSSs (Figure 4A-B, S7). Unlike A485, no difference was observed in the level of K27ac at active TSSs following inhibition of transcription initiation in the presence of TSA (Figure 4B). This suggested that the increase in K27ac at active TSSs was caused by a loss of HDAC activity following inhibition of transcription initiation.

TSA had no effect on the overall level of H2AZ but did result in an overall increase in H3K4me3 (Figure 4A, C-D, S7). However, H3K4me3 levels remained unaffected by triptolide in TSA treated cells (Figure 4C). Conversely, inhibition of transcription initiation still enhanced H2AZ incorporation after TSA treatment (Figure 4D). Together, this indicated that histone acetylation and H2AZ incorporation were independently regulated at active TSSs and both processes were enhanced by inhibition of transcription initiation.

### The relationship between transcription and histone modifications

In this study, we show that inhibition of transcription initiation causes an increase in histone acetylation and H2AZ incorporation at active TSSs. Furthermore, blocking transcription initiation does not prevent *de novo* histone acetylation at hormone induced TSSs and enhancers. We thus conclude that the placement of these active chromatin marks is not specifically dependent on transcription.

While inhibition of p300/CBP HAT activity reduced the overall levels of acetylation at TSSs, it did not prevent the triptolide-induced increase in acetylation. This indicated that the gained acetylation at TSSs was not dependent on the deposition of new acetylation by HATs.

Conversely, HDAC inhibition increased the overall levels of TSS acetylation, and no further increased in histone acetylation was observed in combined TSA and triptolide treatment. As the gained acetylation was fully masked by HDAC inhibition, this suggested that inhibition of transcription initiation also resulted in a loss of HDAC activity at active TSSs.

H2AZ incorporation occurred independently of both transcription and histone acetylation, as neither the overall H2AZ levels or the effect of triptolide on H2AZ incorporation were altered by HAT or HDAC inhibition. Thus, inhibition of transcription initiation was independently associated with both a loss of HDAC activity and the retention of H2AZ. HDACs and the TIP60/P400, SRCAP, and INO80 complexes that regulate H2AZ incorporation indeed localize to active TSSs(*20-23*). Our findings suggest that their activity is recruited or promoted by the initiation of transcription. Intriguingly, live imaging studies have shown that acetylation precedes initiation and that initiated RNAP2 and K27ac modified histones occupy separate chromatin regions(*24, 25*). These observations support a model in which the presence of acetylation and H2AZ poises TSSs for initiation, upon which these marks are actively removed from the chromatin.

Auxin-inducible degradation of RNAP2 alone reproduced the effect of triptolide on acetylation and H2AZ levels at TSSs. However, while similar reduction of RPB1 levels were achieved with 2 hours of auxin or triptolide in HCT116-mAID-POLR2A cells, degradation alone yielded a smaller increase in acetylation and H2AZ levels than observed following triptolide treatment. As the remaining RNAP2 in auxin-treated cells may still initiate transcription, this suggests that the process of transcription initiation recruits or facilitates the activity of HDACs and the complexes that govern H2AZ incorporation at active TSSs. Indeed, RNAP2 generates unwraps DNA from nucleosomes and generates hexasomes, providing more enzymatically favorable substrates for HDACs and H2AZ remodelers(*10, 26-28*).

At active TSSs, histone deacetylation and H2AZ removal are transcriptionally repressive or inhibitory changes to the chromatin landscape. As such, their association with transcription initiation appears counterintuitive. However, these changes may represent a transcriptionally coupled mechanism to govern gene activity. Histone acetylation influences transcriptional bursting dynamics, which can be measured by how often an individual allele is actively transcribed (burst frequency) and/or the number of mRNA produced when an allele is activated (burst size). Targeted approaches using CrisprA demonstrate that directing acetylation to specific promoters increases the size and frequency of transcriptional bursts global inhibition of HDACs also leads to increases in burst size (*29*). Thus, transcription associated HDAC activity may function in the native context to limit bursting dynamics.

In contrast to the changes in histone acetylation and H2AZ incorporation, the levels of H3K4me3 were unaffected by inhibition of transcription initiation. Furthermore, inhibition of HAT activity did not alter the levels of H3K4me3. This supports findings that suggest that H3K4me3 precedes K27ac in the hierarchy of TSS modifications and functions a bookmark for active TSSs (*30*). Indeed, *de novo* induction of H3K4me3 by CrisprA also induces K27ac(*31*). Additionally, while H3K4me3 and K27ac can both induce transcriptional activation when installed *de novo* at a promoter, the effect of *de novo* H3K4me3 was dramatically reduced by inhibition of HAT activity(*31*). Therefore, our findings support a model suggesting that H3K4me3 marks TSSs that can be activated, and subsequent histone acetylation promotes initiation of transcription.

Surprisingly, the effect of blocking transcription initiation on histone acetylation levels was limited to active TSSs and did not have a robust effect on histone acetylation at active enhancers.

RNAP2 and general transcription factors including TFIID and TFIIH are recruited to both TSSs and enhancers (*32*). This suggests that an underlying difference between TSSs and enhancers alters the ability of RNAP2 to recruit or direct the activity of histone modifying proteins. The methylation status of H3K4 may play a role in this difference. While mono-, di-, and tri-methylation of H3K4 are observed at both enhancers and TSSs, the ratio of H3K4me1:H3K4me3 is increased at enhancers(*33*). However, the differences between enhancers and TSSs are not definitive, and it can be difficult to distinguish enhancers and TSSs based solely on their chromatin landscapes. Further examination of the determinants governing the effect of transcription on histone modifications at TSSs may provide greater distinction between TSSs and enhancers.

If the role of histone acetylation at TSSs is to promote bursting and/or initiation of transcription, then genome-wide hyperacetylation of TSSs would likely cause global alterations in the cellular transcriptional profile. The hyperacetylation observed after triptolide treatment likely represents a greater proportion of alleles with acetylated TSSs. After recovering from triptolide treatment, RNAP2 is the probable rate-limiting factor in the reestablishment of the transcriptome. As RNAP2 levels recover, the increased number of acetylated and initiation competent TSSs may result in a flattening of the global transcriptional profile, with enhanced expression of low burst size genes and dampened expression of high burst size genes. Indeed, several studies have demonstrated that HDAC inhibition has complex effects on gene expression and often broadly disrupts the transcriptome. A striking example of this was observed during zygotic genome activation (ZGA) in mouse embryos, the process during which transcription first occurs in the embryonic genome. HDAC inhibition prior to ZGA resulted in global alteration of the transcriptome once ZGA began, with significant changes in expression observed at ∼3,000 genes(*34*). Additionally, inhibition of P300/CBP HAT activity greatly disrupted the transcriptome during post-mitotic genome activation, indicating a requirement for histone acetylation in reestablishing the transcriptome following (*30*). As such, histone acetylation levels appear to be both determined by and a determinant of active transcription.

## Acknowledgements

We are grateful to the members of the Archer laboratory, the NIEHS Epigenetics and Stem Cell Biology Laboratory, and the NIEHS Integrative Bioinformatics Group for ongoing support, advice, and constructive criticism. We also thank G. Solomon, J. Malphurs, N. Reeves, and X. Xu of the NIEHS Epigenomics Core Laboratory for next-generation sequencing expertise.

Finally, we thank P. Wade, J. Watts, G. Muse, and D. Theofilatos for critical review of the manuscript and data.

## Author Contributions

Project Design: TKA. Conceptualization: J.A.H, K.W.T, and T.K.A. Data curation: J.A.H. Formal Analysis: J.A.H. Funding acquisition: T.K.A. Investigation: K.W.T. and J.A.H. Methodology: J.A.H and K.W.T. Supervision: T.K.A. Validation: J.A.H. and K.W.T. Visualization: J.A.H. Writing – original draft: J.A.H. Writing – review & editing: J.A.H., K.W.T., and T.K.A.

## Data and materials availability

All data needed to evaluate the conclusions in the paper are present in the paper and/or the Supplementary Materials. All sequencing data generated for this study have been deposited at GEO: GSE277183.

**Figure S1:**
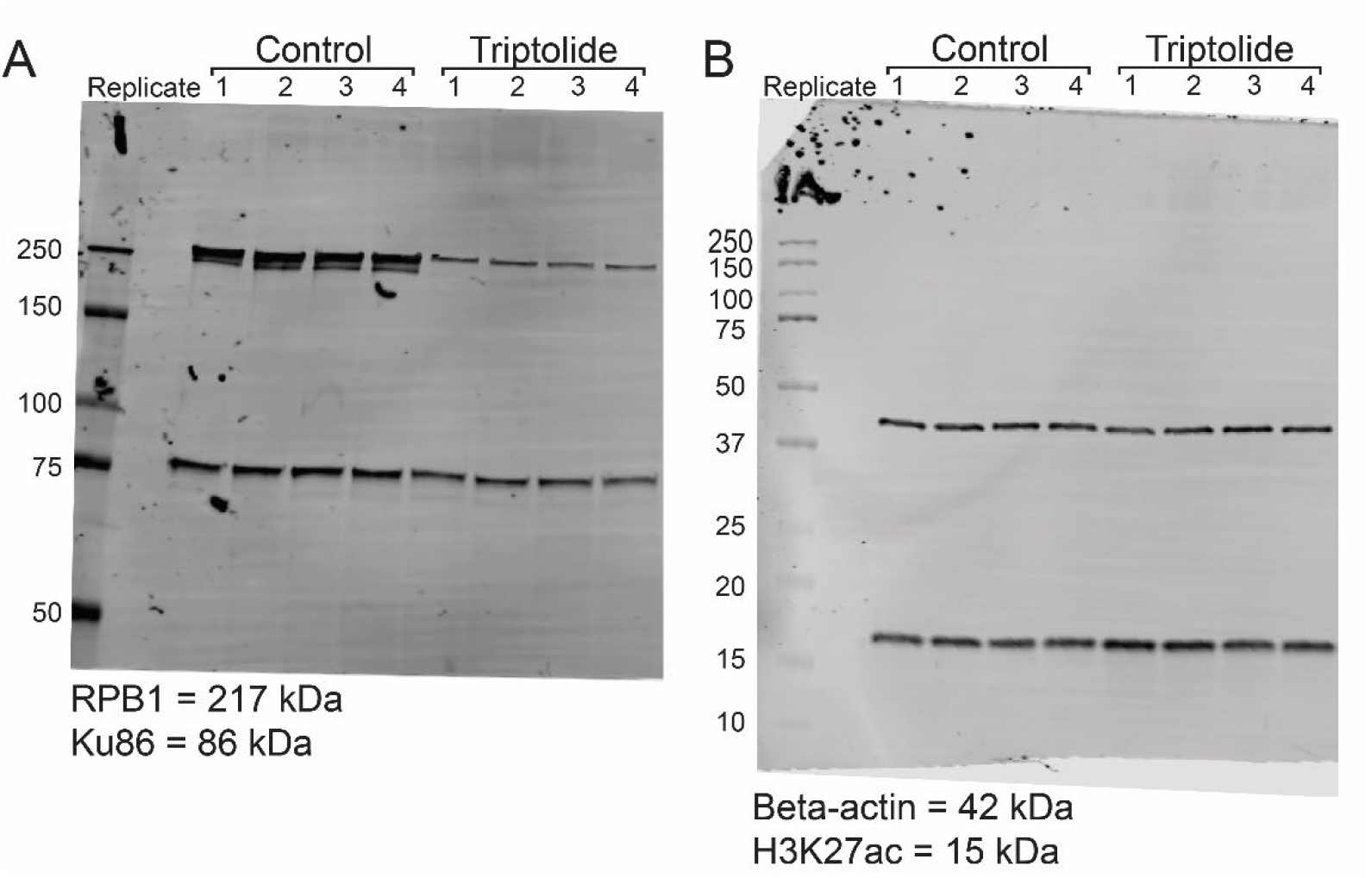
Uncropped western blot images corresponding to Figure 1D.

**Figure S2:**
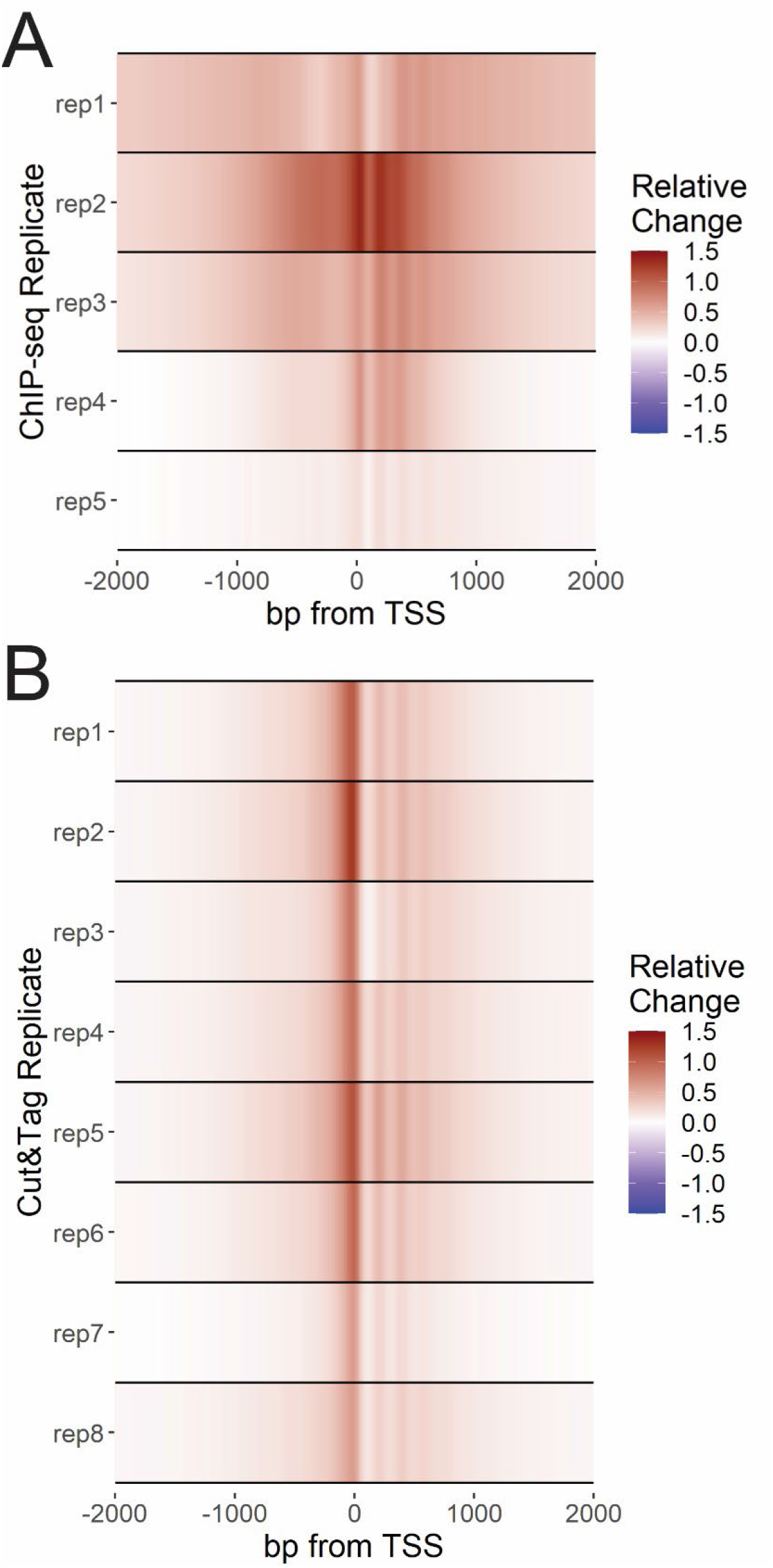
A) Heatmaps depicting relative change ((Triptolide – control) / max(control)) in K27ac ChIP-seq signal across 5 independent biological replicates. B) Heatmaps depicting relative change ((Triptolide – control) / max(control)) in K27ac Cut&Tag signal across 8 independent biological replicates.

**Figure S3:**
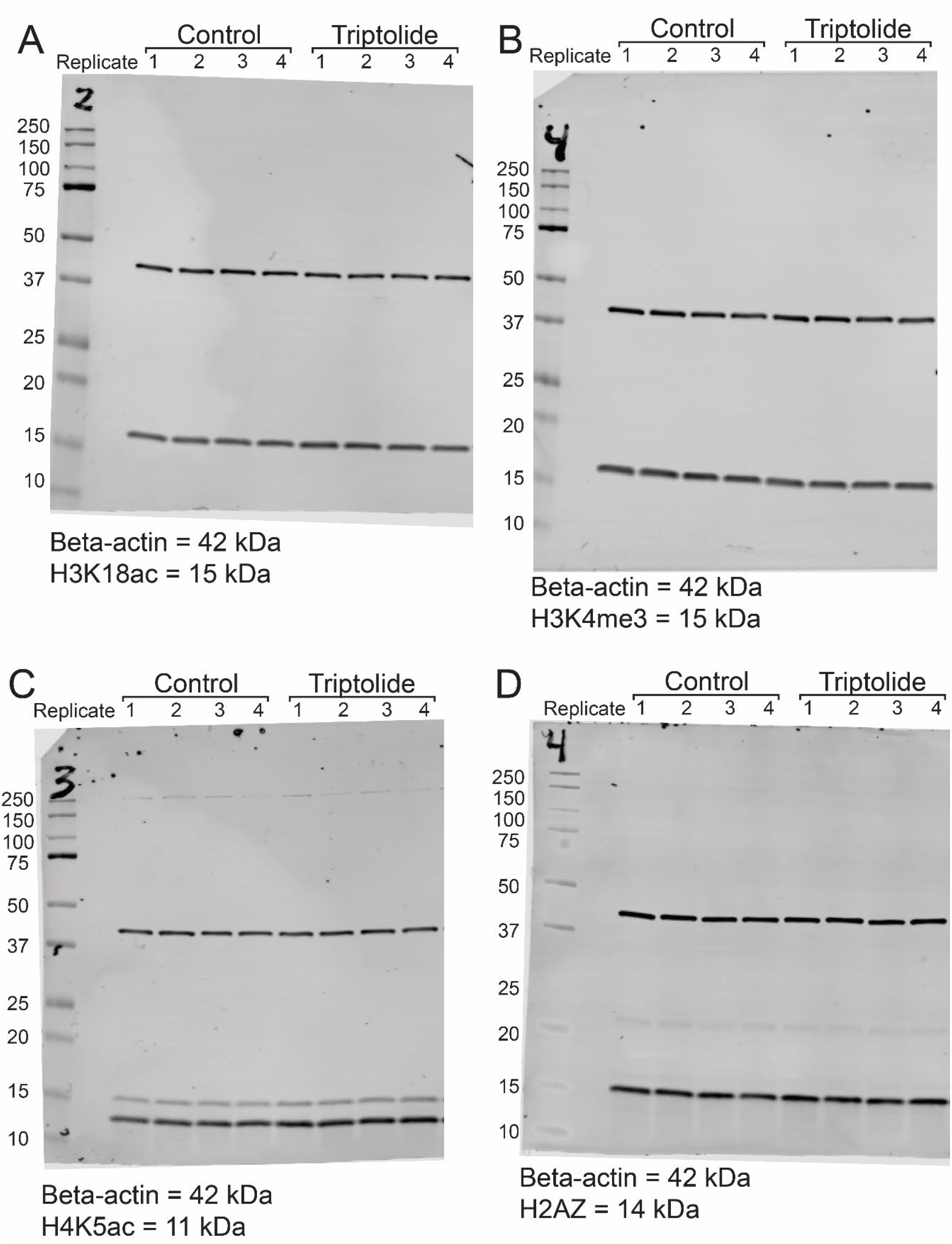
Uncropped western blot images corresponding to Figure 2.

**Figure S4:**
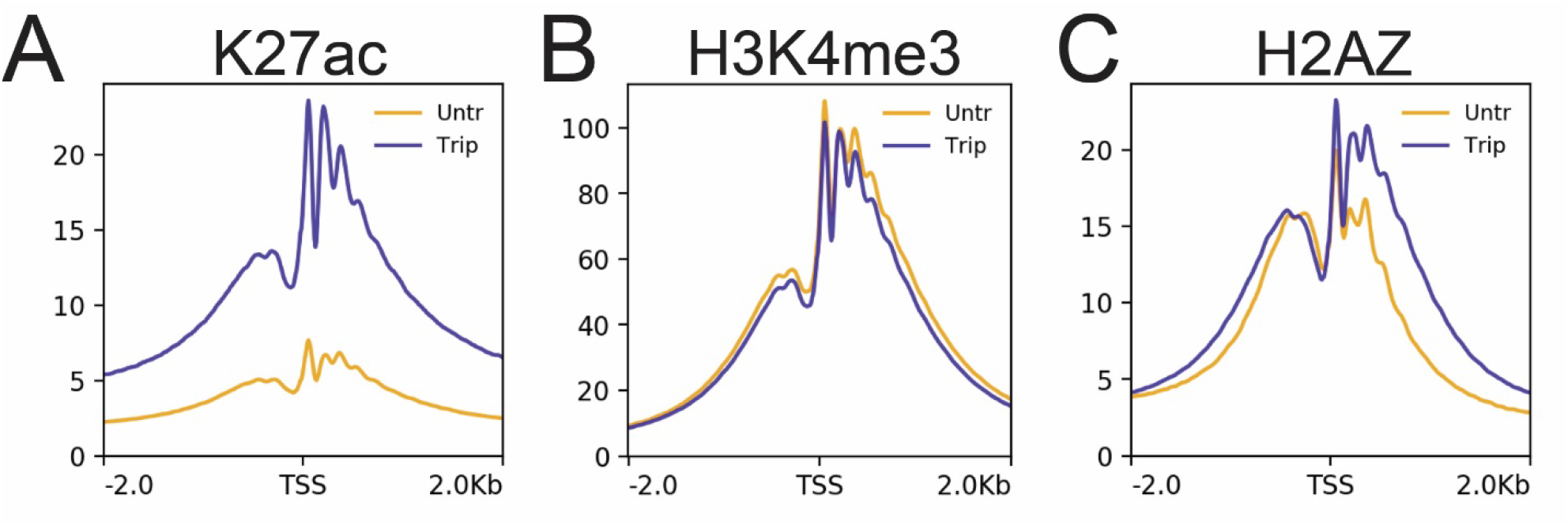
Meta-profiles of ChIP-seq signal +/-2 hours triptolide over Refseq TSSs in K562 cells processed with mild fixation and MNase fragmentation protocol.

**Figure S5:**
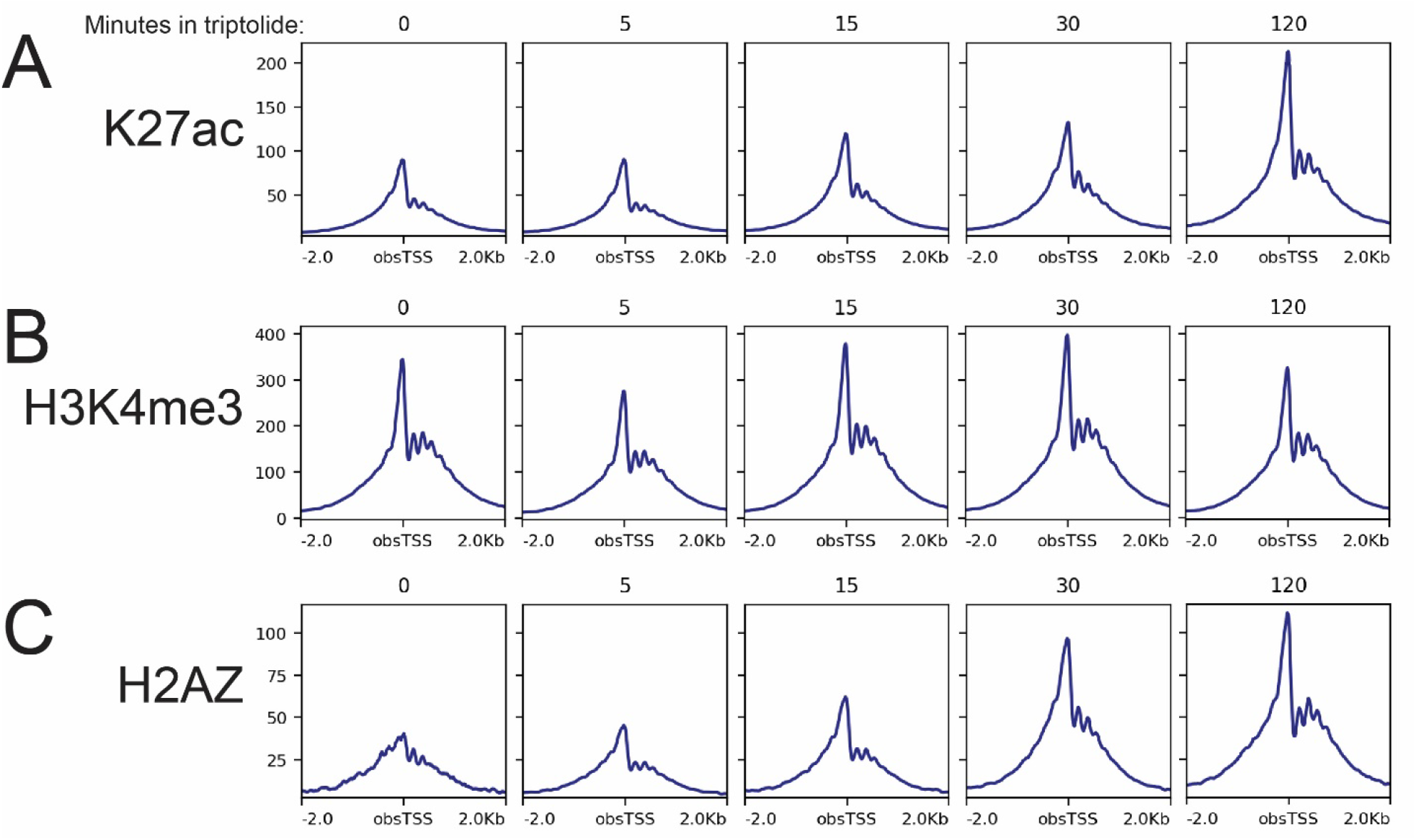
Meta-profiles of Cut&Tag signal +/-indicated minutes of triptolide treatment over active TSSs in A1-2 cells.

**Figure S6:**
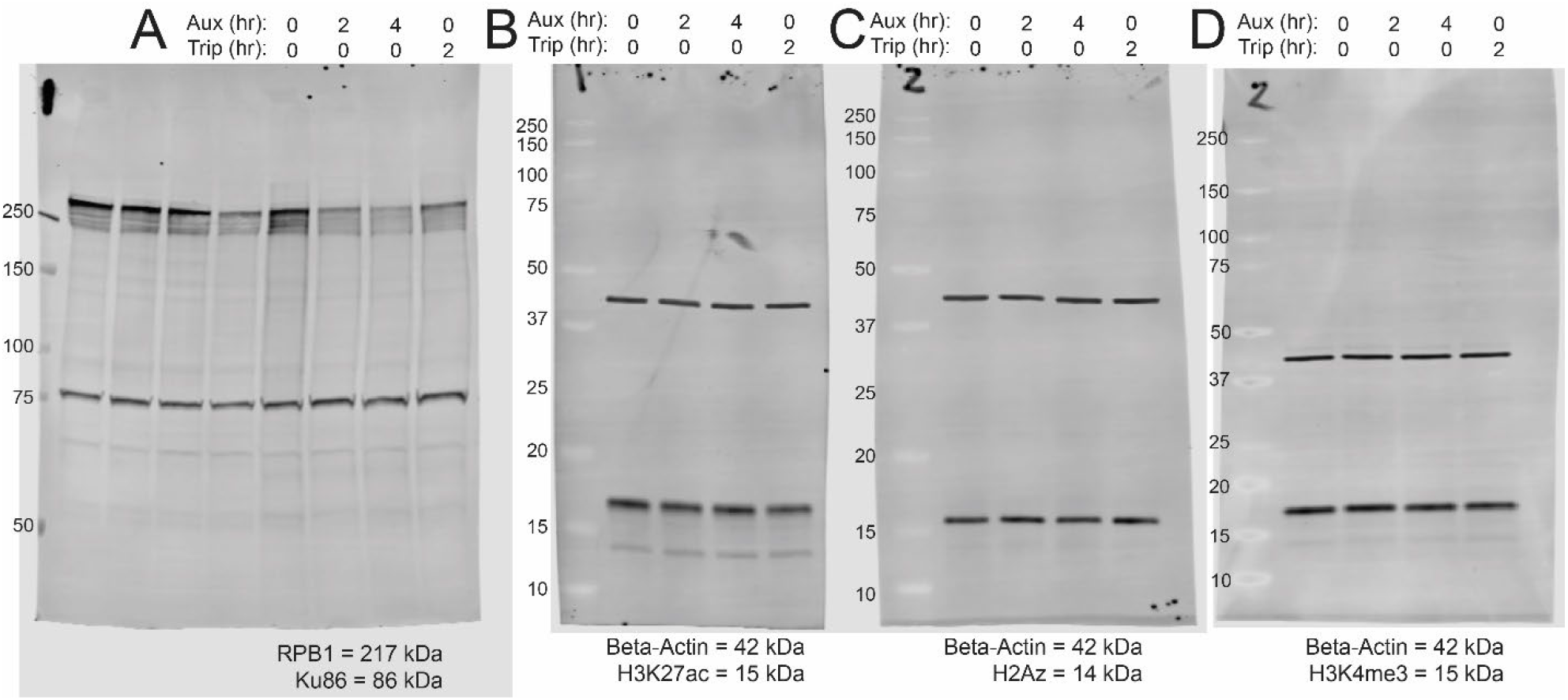
Uncropped western blot images corresponding to Figure 3E.

**Figure S7:**
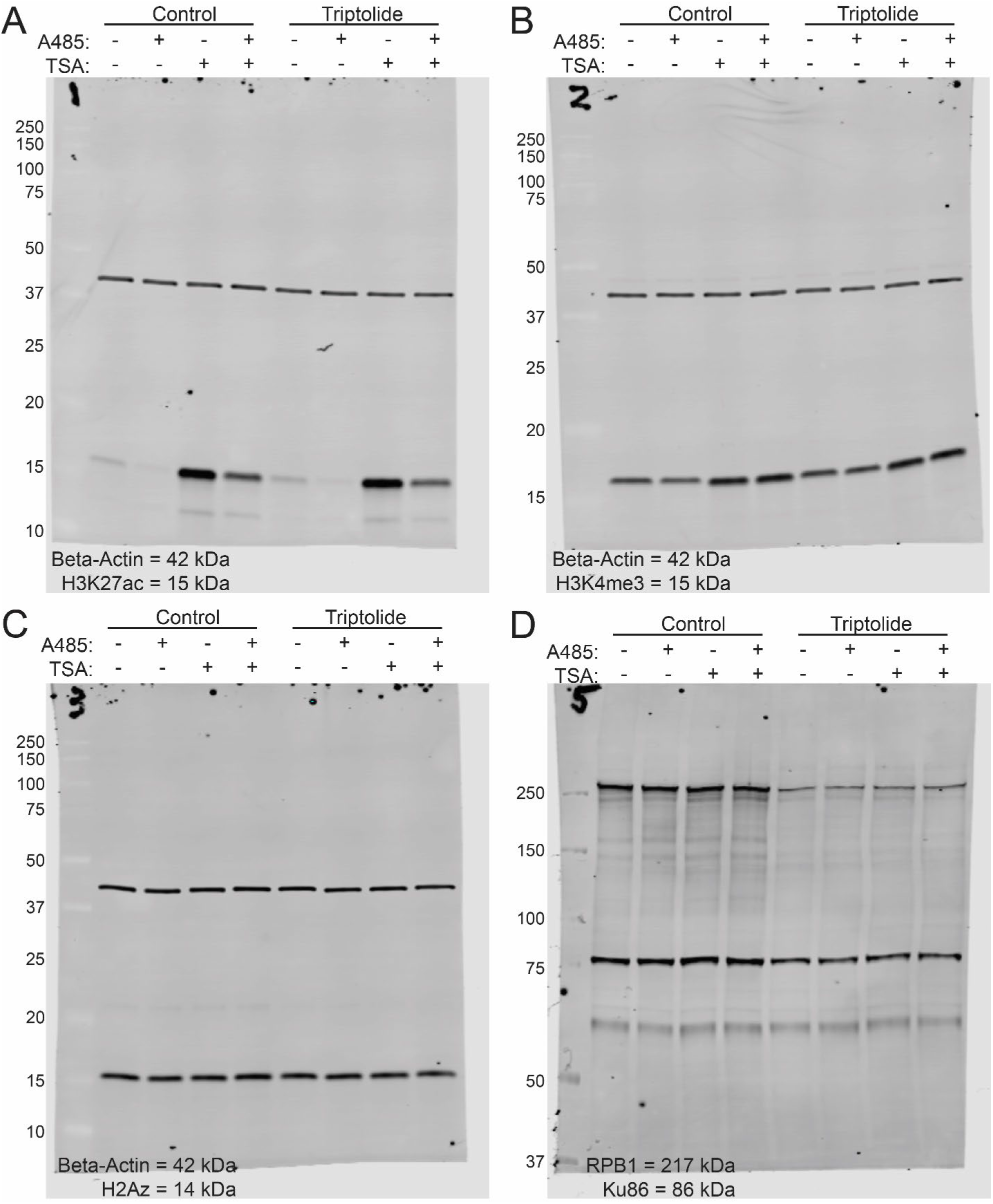
Uncropped western blot images corresponding to Figure 4A.

