## Supplementary material for "RNA Polymerase II coordinates histone deacetylation at active promoters": Methods and Supplemental Figures

Materials and Methods

**Cell lines and treatments.**

Human T47D A1-2 (Archer et al., 1994) and MCF-7 cell were grown in MEM (Gibco 10370) supplemented with 10% fetal bovine serum (Gibco 26140-079), GlutaMAX (Gibco 3505), HEPES (Sigma H0887), and Penicillin/Streptomycin (Sigma P0781). HeLa, SW-13, U2OS, primary human fibroblast and mouse C127 cells were grown DMEM (Gibco 11965-092) supplemented with 10% fetal bovine serum (Gibco 26140-079), GlutaMAX (Gibco 3505), HEPES (Sigma H0887), and Penicillin/Streptomycin (Sigma P0781). K562 cells were grown in suspension in RPMI Medium 1640 (Gibco 11875-093) supplemented with 10% fetal bovine serum (Gibco 26140-079), GlutaMAX (Gibco 3505), HEPES (Sigma H0887), and Penicillin/Streptomycin (Sigma P0781).

All cells were cultured in a humidified incubator at 37°C with 5% CO2 in the indicated growth medium with media changed every 2-3 days. Cells were seeded in new tissue culture treated dishes when 80% confluency reached.

Cells were treated for the indicated time individually, or in combination, with 100 nM Dexamethasone (Sigma D4902), 0.5 uM Triptolide (Tocris 3253), 0.5 uM Flavopiridol (Tocris 3094), 0.5 uM A485 (Tocris 6387), 0.1 uM Trichostatin A (Sigma T8552), 10 ug/ml Doxycycline (Sigma D9891), 1 uM 5-Ph-IAA (AID) ligand (Sigma SML3574), or vehicle (equal volume EtOH or DMSO).

**Immunoblot analysis.**

Whole cell and nuclear protein extracts were prepared from cell or nuclei aliquots taken after indicated treatment and prior to down-stream assay analysis. Compact cell or nuclei pellets were resuspended in RIPA-HS buffer (25 mM Tris HCl pH 7.6, 500 mM NaCl, 1% NP-40, 1% sodium deoxycholate, 0.1% SDS, 100 uM PMSF, and 1x cOmplete EDTA-free protease inhibitors (Roche)), incubated on ice for 15 min, then subjected to sonication using PIXUL Multi-Sample Sonicator (Active Motif) at 12C using the following parameters: Pulse [N]: 50; PRF [kHz]: 1.00; Process Time: 60 sec; Burst Rate [Hz]: 20.00. After sonication, samples were centrifugated at 14,000g for 15 min and protein extract collect as supernatant. Protein concentration determined by Bradford assay. Whole cell (50 ug) or nuclear (5 ug) were resolved using SDS-PAGE, transferred to low fluorescence PVDF membrane (Bio-rad), and blocked in TBS Blocking Buffer (Li-Cor). Immunoblot analysis was performed as previously described(*35*) using the indicated antibodies as described in Table S1 with a direct near-infrared fluorescence detection system (Li-cor Odyssey CLx Imaging System).

**ChIP-seq.**

Cells were fixed in PBS with 1% methanol-free formaldehyde (ThermoFisher Scientific 28906) at 37C for 10 min and quenched with glycine (0.125M) for 5 min at room temperature. After glycine quench, cells were washed three times and scraped into ice-cold PBS and pelleted by centrifugations at 500g for 5 min. Compact cell pellets were resuspended in MNase swelling buffer (25 mM HEPES pH 7.9, 1.5 mM MgCl2, 10 mM KCl, 0.1% NP-40, 0.5 mM PMSF, and 1x cOmplete EDTA-free protease inhibitors (PIC)) (Roche), incubated on ice for 10 min then subjected to Dounce homogenization using 20-strokes with “tight” pestle (Duran Wheaton Kimble, 357542). Nuclei were sedimented through digestion buffer (15mM Hepes pH 7.9, 60mM KCl, 15mM NaCl, 0.32M Sucrose, 0.5 mM PMSF, and PIC) by centrifugation for 930g for 7 min at 4C. Nuclei pellets were fully resuspended in digestion buffer containing 3.3 mM CaCl2 and digested with MNase (0.75U/1x10ˆ6 cells, Worthington, cat#) at 37C for 15min. MNase fragmentation was stopped by addition of 10 mM EGTA and incubation for 5 min on ice. Digested nuclei were centrifuged at 930g for 7 min at 4C and compact nuclei pellets resuspended in 20 volumes (v/v) ChIP Shearing buffer (50 mM HEPES pH7.9, 140 mM NaCl, 1 mM EDTA, 1% Triton X-100, 0.1% Na-deoxycholate, 0.1% SDS containing spermidine, PIC and PMSF) and incubated on ice for 10 min. Samples were subjected to sonication using PIXUL Multi-Sample Sonicator (Active Motif) at 12C using the following parameters: Pulse [N]: 50; PRF [kHz]: 1.00; Process Time: 10 min; Burst Rate [Hz]: 20.00. After sonication, fragmented chromatin was recovered in supernatant after centrifugation at 18,000g for 10 min with extent of fragmentation determined by agarose gel electrophorese prior to immunoprecipitation. The concentration of fragmentated chromatin determined by Bradford assay. Fragmentated chromatin was diluted in two-fold in 2xIP buffer (20 mM Tris-HCl pH 8.0, 300 mM NaCl, 2 mM EDTA, 20% Glycerol, 1% Triton X-100, 0.5 mM PMSF and PIC) and immunoprecipitation performed with the indicated antibodies at a ratio of 1 ug antibody per 200 ug chromatin. Equal amounts of Spike-IN antibody and Drosophila chromatin (Active Motif 61686 and 53083) added to each immunoprecipitation for normalization. Immune complexes were captured using protein G dynabeads (Invitrogen 10004D), washed once each with low salt buffer (20 mM Tris-HCl pH 8.0, 150 mM NaCl, 2 mM EDTA, 1% Triton X-100, 0.1% SDS), high salt buffer (same as low salt buffer, except 500 mM NaCl), and LiCl buffer (Tris-HCl pH 8.0, 250 mM LiCl, 2 mM EDTA, 1 % NP-40, 1% (wt/vol) sodium deoxycholate), and twice with TE. Immunoprecipitated DNA recovered from beads by incubation in ChIP Elution Buffer (1% SDS, 0.1M Sodium Bicarbonate) for 30 min at room temperature. Eluted DNA was treated with RNaseA (Roche 10109169001) and Proteinase K (Invitrogen #25530015) and purified using Qiagen PCR purification columns.

ChIP-seq in K562 cells with milder cross-linking and fragmentation performed using a SimpleChIP Plus Enzymatic Chromatin IP Kit with Magnetic Beads (Cell Signaling Technologies #9005S) according to manufacturer’s protocol.

ChIP-seq libraries were generated using the xGen ssDNA & Low-Input DNA Library Prep Kit and xGen CDI Primers (IDT 10009817 and 10009815) according to manufacturer’s protocol and sequenced on the Illumina NextSeq platforms.

**ChIP-seq analysis.**

Reads were trimmed using Cutadapt (*36*) and aligned to both hg19 and dm6 genomes using bowtie2 (*37*). Aligned reads were deduplicated with picardtools (http://broadinstitute.github.io/picard). Scale factors were calculated for each sample by dividing the minimum number of dm6 reads for a given antibody by number of dm6 reads for each sample using the same antibody. Scaled and normalized coverage files were generated using deeptools (*38*) with calculated scale factors and rpkm normalization. Meta-profiles and heatmaps were generated using deeptools and ggplot2.

**CUT & TAG**

Cells were treated, harvested, and subjected to CUT&TAG analysis using a modified published protocol(*39*). Trypsinized cells were washed once with PBS and incubated with NE nuclear extraction buffer (20 mM HEPES-KOH, pH 7.9, 10 mM KCl, 0.1% Triton X-100, 20% Glycerol, 0.5 mM Spermidine, 100 uM PMSF, and 1x EDTA-free protease inhibitors (Roche cOmplete) for 30 min on ice followed by Dounce homogenization using 20-strokes from Dounce Tissue Grinder (Wheaton 357542) tight pestle. Following homogenization, samples were centrifuged for 3 min at 600g, nuclei pellet resuspended in NE buffer supplemented with 0.1 % formaldehyde and incubated at room temperature for 2 minutes followed by quench with 0.125 M glycine. Following formaldehyde cross-link, nuclei were re-centrifuged, resuspended in fresh NE buffer and counted with trypan blue dye using Bio-Rad TC20 Automated Cell Counter.

After isolation, 150,000 total nuclei [135,000 nuclei from experimental sample with 15,000 mouse spike-in nuclei] were incubated with activated concanavalin A-coated (ConA) magnetic beads (EpiCypher) in 0.2 ml thin-walled tubes for 10 min at room temperature. Nuclei-bound ConA beads were resuspended in antibody incubation buffer (20 mM HEPES, pH 7.5, 150 mM NaCl, 0.01% Digitonin, 2 mM EDTA, 0.5 mM Spermidine, 100 uM PMSF, and 1x EDTA-free protease inhibitor), 0.75 ug indicated primary antibody (see Table S2) added and samples incubated overnight at 4C with gently rotation. Primary antibody removed from ConA bead solution by placing samples on magnetic stand until clear followed by supernatant removal. Samples were incubated with 0.75 ug species specific secondary antibody for 30 min at room temperature with gentle rotation followed by two washes with Digitonin150 wash buffer (20 mM HEPES, pH 7.5, 150 mM NaCl, 0.01% Digitonin, 0.5 mM Spermidine, 100 uM PMSF, and 1x EDTA-free protease inhibitor) then incubation with pAG-Tn5 in Digitonin300-wash buffer (20 mM HEPES, pH 7.5, 300 mM NaCl, 0.01% Digitonin, 0.5 mM Spermidine, 100 uM PMSF, and 1x EDTA-free protease inhibitor) for 1h at room temperature.

After pAG-Tn5 incubation, bound ConA beads were washed twice with Digitonin300-wash buffer then resuspended in Tagmentation buffer (Digitonin300 wash buffer containing 10 mM MgCl2) and incubated for 1h at 37C. Beads were then pelleted, resuspended in TAPS buffer (10 mM TAPS pH 8.5, 0.2 mM EDTA). TAPS buffer removed and beads carefully resuspended in SDS Release buffer (10 mM TAPS pH 8.5, 0.1% SDS) and incubated for 1 h at 58 C. Following incubation, SDS Quench buffer (0.67% Triton X-100) was added to each sample along with 2 μl of a universal i5 and a uniquely barcoded i7 primers (from 10 μM stocks). Equal volume (25 ul) NEBNext High-Fidelity 2x PCR Master mix (M0541, New England Biolabs) was added to each sample and mixed by gentle pipetting. Samples subjected to DNA amplification using a thermocycler and the CUT&TAG-specific PCR cycling parameters: 58C for 5 min; 72C for 5 min; 98C for 45 sec; with 19 cycles of 98C for 15 sec and 60C for 10 sec; final extension at 72 °C for 1 min; and hold at 4C. DNA clean-up performed by adding 1.3x AMPure XP beads (A63880, Beckman Coulter) to each sample, allowing libraries to incubated with beads for 10 min at room temperature, followed by two washes with 80% ethanol, then elution in 15 μl of 10 mM Tris pH 8.0. While on magnetic stand, supernatant was carefully taken and transferred to a fresh tube. Library size distribution determined by Agilent TapeStation 4200 (Agilent Technologies) and libraries mixed to yield equal representation prior to paired-end (2 x 50 bp) Illumina sequencing by NIEHS The Epigenomics and DNA Sequencing Core Facility.

**CUT & TAG analysis.**

Reads were trimmed using Cutadapt (*36*) and aligned to both hg19 and mm10 genomes using bowtie2(*37*). Aligned reads were deduplicated with picardtools. Deduplicated read counts were used to calculate the human:mouse ratio, which was then used to calculate scaling factors for samples performed side by side with the same antibody. The scale factor was calculated by dividing the human:mouse read ratio for each sample using a given antibody by the maximum human:mouse read ratio from samples using the same antibody. Scaled and normalized coverage files were generated using deeptools (*38*) with calculated scale factors and rpkm normalization. Meta-profiles and heatmaps were generated using deeptools and ggplot2.

**ATAC-seq**

Two replicates were used for each treatment condition. Isolated cells were treated in 10 mM PIPES, pH 6.8, 100 mM NaCl, 300 mM sucrose, 3 mM MgCl2, and 0.1% Triton X-100 and then treated with the Illumina Tagment DNA TDE1 Enzyme and Buffer Kits for 30 min followed by mixing every 10 min. Isolation of libraries and amplification was performed as described (*40*). Libraries were sequenced at the NIEHS Epigenomics Core Facility on a NovaSeq instrument, and reads were trimmed with default parameters.

35.	N. Dietrich, K. Trotter, J. M. Ward, T. K. Archer, BRG1 HSA domain interactions with BCL7 proteins are critical for remodeling and gene expression. *Life Science Alliance* **6**, e202201770 (2023).

36.	M. Martin, Cutadapt removes adapter sequences from high-throughput sequencing reads. *EMBnet.journal; Vol 17, No 1: Next Generation Sequencing Data AnalysisDO - 10.14806/ej.17.1.200*, (2011).

37.	B. Langmead, S. L. Salzberg, Fast gapped-read alignment with Bowtie 2. *Nature Methods* **9**, 357-359 (2012).

38.	F. Ramírez *et al.*, deepTools2: a next generation web server for deep-sequencing data analysis. *Nucleic Acids Research* **44**, W160-W165 (2016).

39.	H. S. Kaya-Okur *et al.*, CUT&Tag for efficient epigenomic profiling of small samples and single cells. *Nature Communications* **10**, 1930 (2019).

40.	J. D. Buenrostro, B. Wu, H. Y. Chang, W. J. Greenleaf, ATAC-seq: A Method for Assaying Chromatin Accessibility Genome-Wide. *Current Protocols in Molecular Biology* **109**, 21.29.21-21.29.29 (2015).

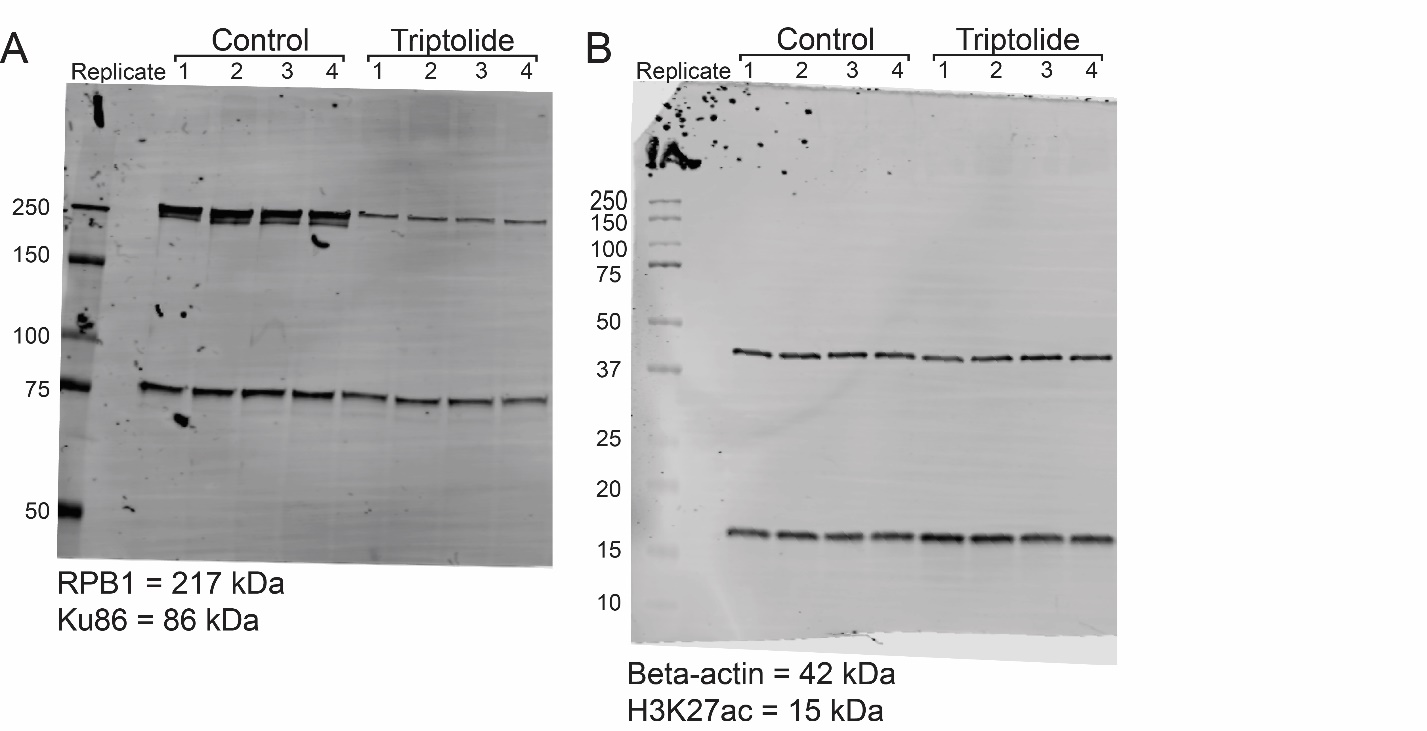

**Figure S1:** Uncropped western blot images corresponding to Figure 1D.

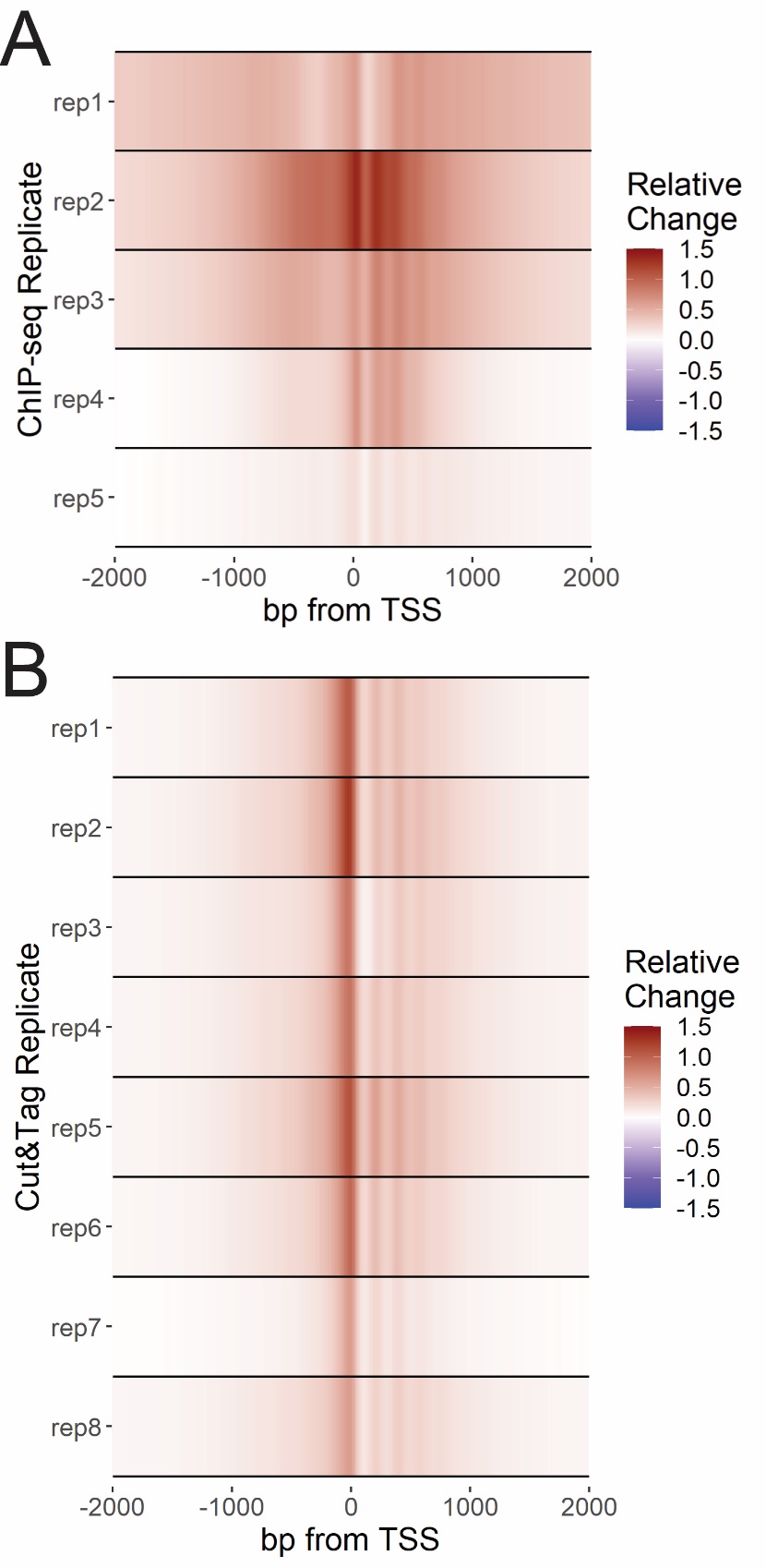

**Figure S2:** A) Heatmaps depicting relative change ((Triptolide – control) / max(control)) in K27ac ChIP-seq signal across 5 independent biological replicates. B) Heatmaps depicting relative change ((Triptolide – control) / max(control)) in K27ac Cut&Tag signal across 8 independent biological replicates.

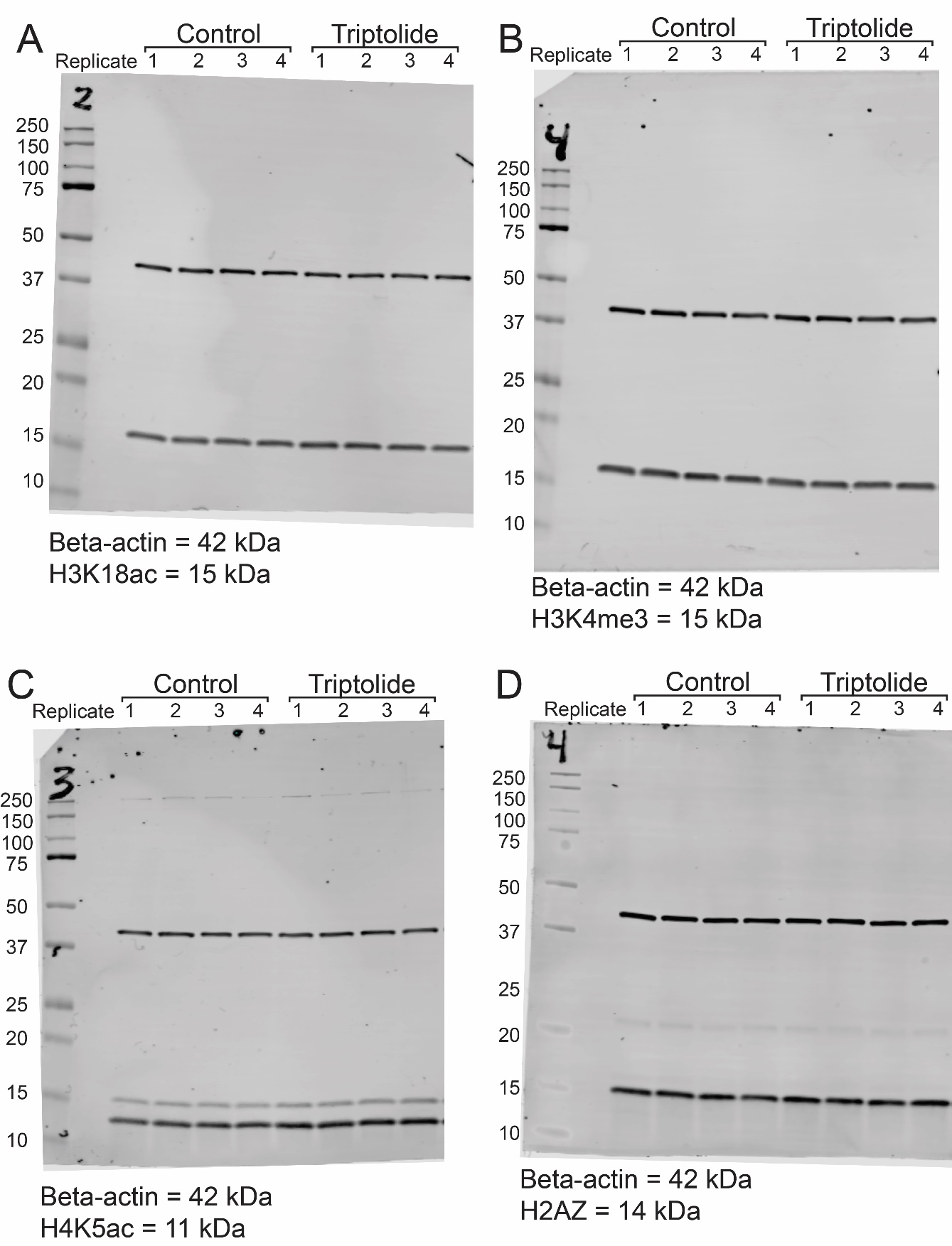

**Figure S3:** Uncropped western blot images corresponding to Figure 2.

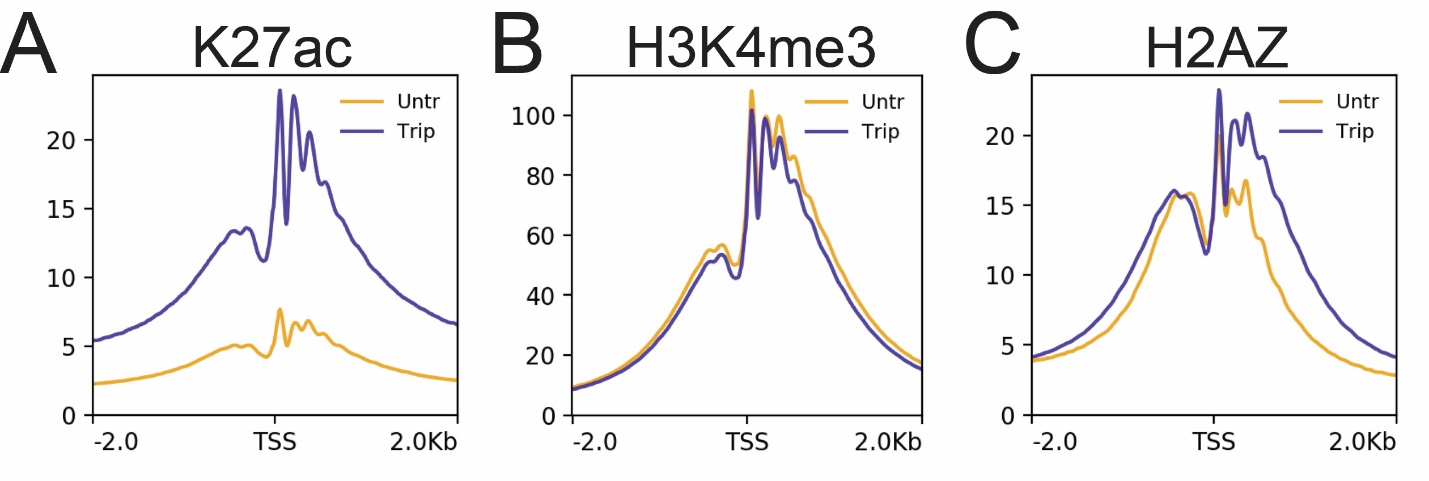

**Figure S4:** Meta-profiles of ChIP-seq signal +/- 2 hours triptolide over Refseq TSSs in K562 cells processed with mild fixation and MNase fragmentation protocol.

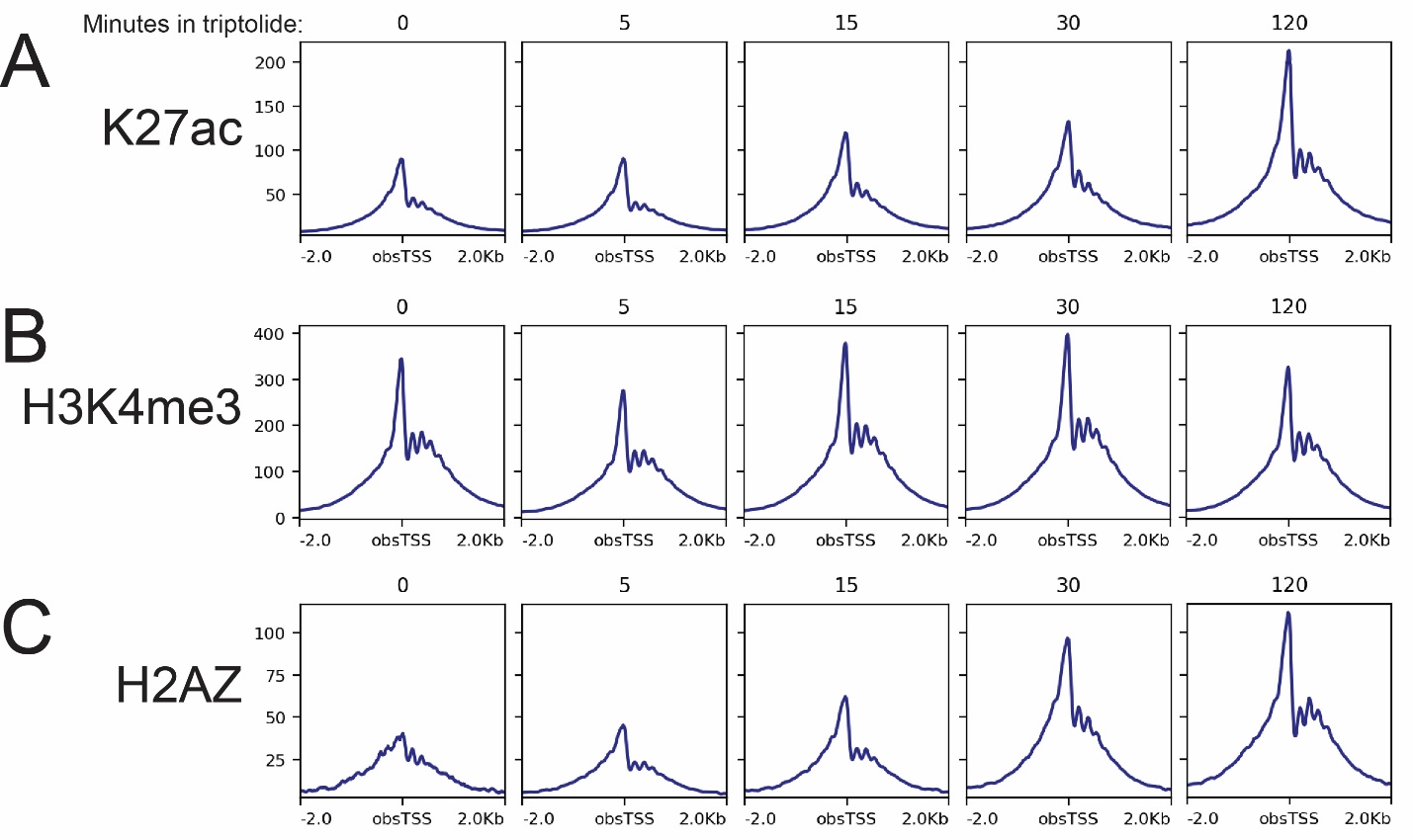

**Figure S5:** Meta-profiles of Cut&Tag signal +/- indicated minutes of triptolide treatment over active TSSs in A1-2 cells.

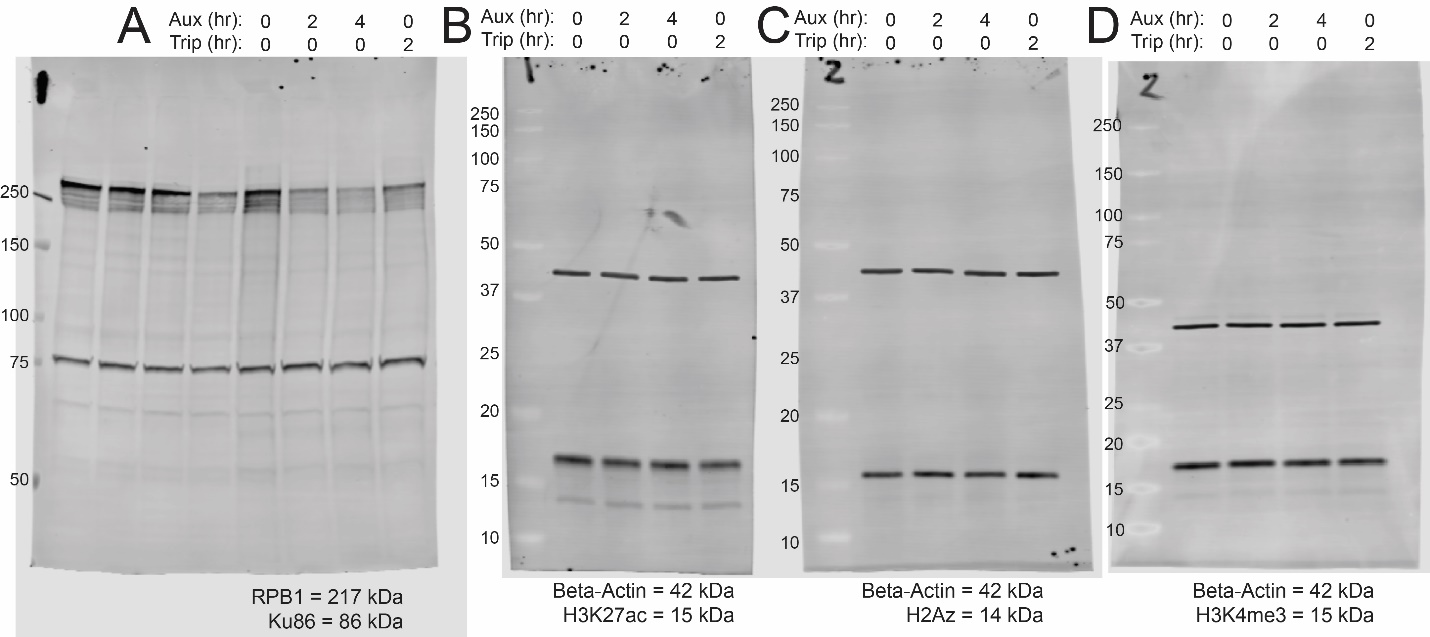

**Figure S6:** Uncropped western blot images corresponding to Figure 3E.

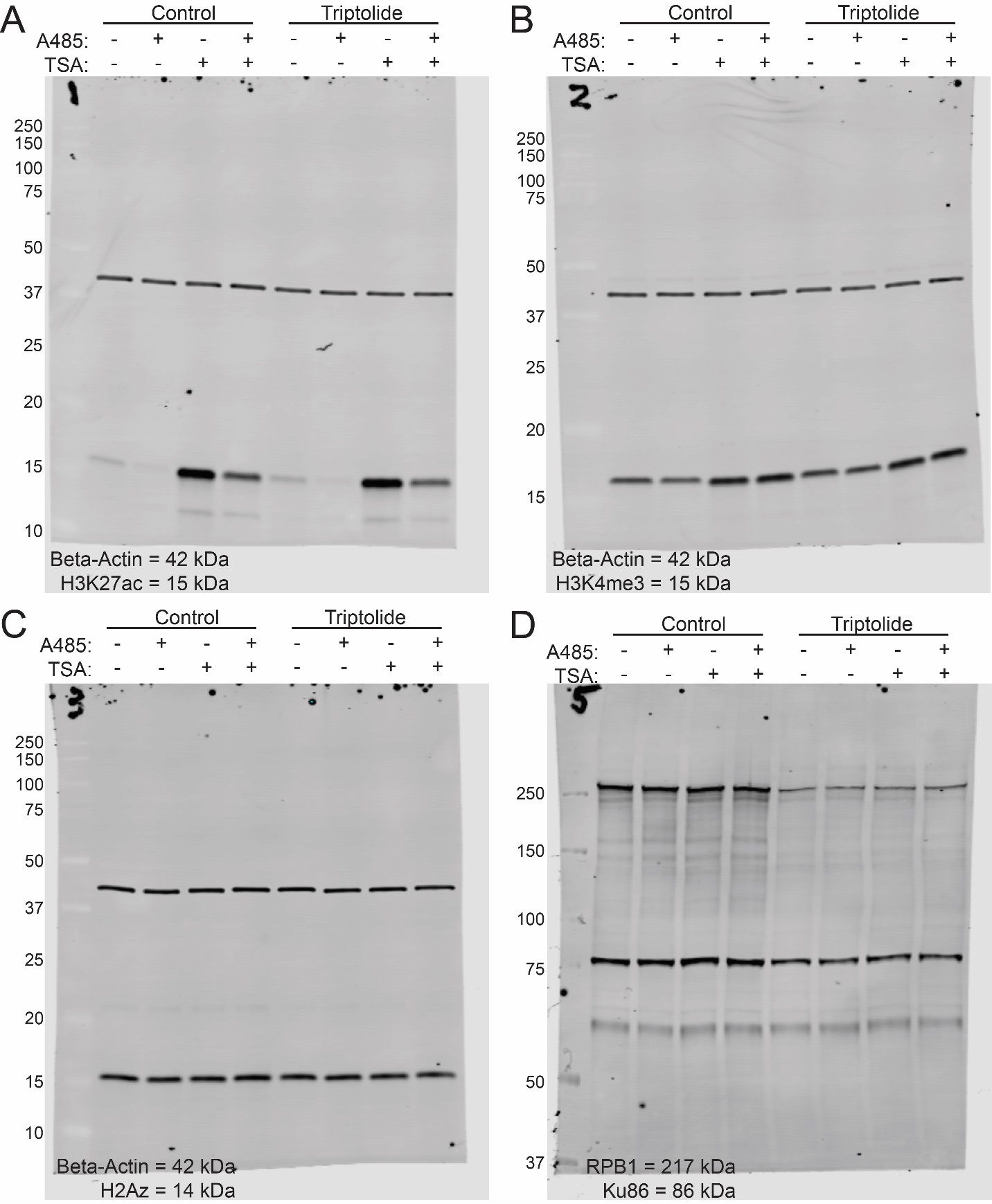

**Figure S7:** Uncropped western blot images corresponding to Figure 4A.

**Table S1: Immunoblot antibodies**

| **Target:** | **Cat. Num:** | **Vendor:** | **RRID:** | **Dilution:** |
| --- | --- | --- | --- | --- |
| RNAP2 | 39097 | Active Motif | AB_2732926 | 0.2 ug/ml |
| Ku86-S10B1 | sc-56136 | Santa Cruz Biotechnology | AB_794204 | 0.2 ug/ml |
| H3K27ac | ab4729 | Abcam | AB_2118291 | 0.5 ug/ul |
| β-Actin | A1978 | Sigma-Aldrich | AB_476692 | 0.5 ug/ml |
| H2A.Z | 39943 | Active Motif | AB_2793401 | 0.5 ug/ml |
| H3K4me3 | C15410003 | Diagenode | AB_2924768 | 0.5 ug/ml |
| H3K18ac | C15410139 | Diagenode | AB_2713907 | 0.5 ug/ml |
| H4K5ac | C15410025 | Diagenode | AB_3661649 | 0.5 ug/ml |

**Table S2: CUT&TAG and ChIP-seq Antibodies**

| **Target:** | **Cat. Num:** | **Vendor:** | **RRID:** |
| --- | --- | --- | --- |
| H3K27ac | ab4729 | Abcam | AB_2118291 |
| H3K4me3 | C15410003 | Diagenode | AB_2924768 |
| H3K9ac | C15410004 | Diagenode | AB_2713905 |
| H2A.Z | C15410201 | Diagenode | AB_3661648 |
| H3K27me3 | C15410195 | Diagenode | AB_2753161 |
| H3K18ac | C15410139 | Diagenode | AB_2713907 |
| H3K14ac | ab52946 | Abcam | AB_880442 |
| RNAP2 | 39097 | Active Motif | AB_2732926 |
| H4K5ac | C15410025 | Diagenode | AB_3661649 |
| H4K12ac | C15410331 | Diagenode | AB_3661650 |
| H3K18ac | C15410139 | Diagenode | AB_2713907 |
| GR/NR3C1 | NBP2-42221 | Novus | AB_2894721 |
